# Individual differences in behavioral effects of xylazine and opioid-xylazine mixtures in male rats

**DOI:** 10.1101/2025.06.25.661558

**Authors:** Kristen L. Woodhouse, Jun-Xu Li

## Abstract

**Rationale:** Recent work in animals suggests xylazine neither enhances the rewarding effects nor intake of fentanyl. Anecdotal evidence from people who use drugs indicates some individuals prefer fentanyl adulterated with xylazine. Systematic examination of pharmacological interactions between xylazine and opioids is needed to understand the disparate findings between preclinical studies and human reports.

**Objectives:** This study examined behavioral interactions between xylazine and opioids in rats to investigate the pharmacology underlying an emerging trend in drug use.

**Methods:** The sedative, hypothermic, subjective, and respiratory effects of xylazine and opioid-xylazine mixtures were examined in male rats. Locomotor activity was measured in an open field, and body temperature changes were measured with a rectal probe. Rats were trained to discriminate 0.04 mg/kg fentanyl, 0.02 mg/kg fentanyl, or 1.5 mg/kg xylazine from saline and were probed with fentanyl, xylazine, or both to observe whether the drug(s) generalized with the training dose. Whole body plethysmography was used to assess the effects of xylazine on respiration.

**Results:** Xylazine depressed locomotor activity and core body temperature, but considerable variability between subjects was observed. In some subjects, xylazine fully substituted for fentanyl, and prolonged the subjective effects of fentanyl. Doses of 1 and 1.78 mg/kg xylazine only partially generalized to the training dose of 1.5 mg/kg xylazine. Xylazine exacerbated the respiratory depressant effects of opioids, and atipamezole reversed the xylazine enhancement of morphine-induced respiratory depression.

**Conclusions:** Individual differences were observed in multiple behavioral measures following xylazine administration and may recapitulate the divisiveness of xylazine reported in people who use drugs.

## Introduction

Xylazine adulteration of illicit fentanyl has become increasingly prevalent in recent years, particularly in the northeastern United States (Johnson et al. 2021; Kariisa et al. 2023). Use of xylazine as an adulterant initially arose in Puerto Rico nearly 20 years before its surge in detection in Philadelphia (D’Orazio et al. 2023). Anecdotal reports suggest the addition of xylazine, or “tranq”, may increase the duration of fentanyl’s euphoric effects, yet its reception amongst people who use opioids is mixed (Reed et al. 2022; Spadaro et al. 2023). Since xylazine was never approved for human use, little is known about this emerging public health threat. Though xylazine has been shown to increase the lethality of fentanyl overdose in preclinical models (Acosta-Mares et al. 2023; Smith et al. 2023), the extent to which xylazine interferes with the efficacy of naloxone reversal remains poorly understood. Despite a lack of scientific evidence, xylazine has been widely purported to lead to naloxone resistant overdose, and thus there is dire need for first responders to have evidence-based information to respond appropriately and effectively to this growing trend.

For decades, xylazine has been used as a veterinary anesthetic, and the few reported instances of xylazine self-administration have generally involved veterinary practitioners or other professions that work with livestock (Carruthers et al. 1979; Gallanosa et al. 1981; Hoffmann et al. 2001; Poklis et al. 1985; Spoerke et al. 1985). Like the antihypertensive drug clonidine, xylazine is an alpha-2 adrenergic receptor (α2AR) agonist, but recent evidence suggests xylazine is also a kappa-opioid receptor (KOR) agonist (Bedard et al. 2024). Since clonidine is used to treat opioid cessation, and since KOR agonists like nalfurafine have been reported to have abuse-limiting effects in both rodents and nonhuman primates (Freeman et al. 2014; Negus et al. 2008; Zamarripa et al. 2020), that some individuals who use drugs actively seek out fentanyl adulterated with xylazine is perplexing and remains to be explained. While human subjects are divided on the subjective effects of xylazine (Reed et al. 2022; Spadaro et al. 2023), reports of xylazine-induced sedation and incidence of tissue necrosis are widespread. Severe cutaneous lesions and tissue necrosis may develop in people who use xylazine (Pergolizzi et al. 2023; Soderquist et al. 2024; Sun et al. 2024; Wei et al. 2023), and while peripheral vasoconstriction is a suspected explanation, the mechanism by which this occurs is currently unknown. In veterinary and scientific practice, xylazine is routinely administered to rodents, cats, horses, and other mammals without resulting skin lesions – with the exception of rodents administered intramuscular ketamine-xylazine (Smiler et al. 1990) – and it is unclear what underlies this species difference in susceptibility to tissue damage.

Recent preclinical work has focused on evaluating the relationship between xylazine and the reinforcing effects of opioids in substance abuse models, as well as the effects of xylazine on opioid-induced hypoxia. Xylazine pretreatment decreased fentanyl responding in rats trained to self-administer fentanyl, and co-administration of fentanyl and xylazine similarly decreased the number of intravenous infusions relative to fentanyl alone (Khatri et al. 2024). Consistently, co-administration of heroin and xylazine suppressed heroin self-administration in male rats, and decreased motivation to consume heroin in a progressive ratio test (Hochstetler et al. 2025), while fentanyl-xylazine combinations did not significantly alter fentanyl reinforcement in male and female rats that underwent a progressive ratio task (St Onge et al. 2024). Furthermore, while mice developed conditioned place preference following repeated treatment with fentanyl, mice that were treated with fentanyl and xylazine did not develop significant place preference (Acosta-Mares et al. 2023). Altogether these preliminary findings raise questions about the nature of opioid-xylazine interactions, the foremost being why it is that some individuals prefer fentanyl adulterated with xylazine when rodent studies have found that xylazine suppresses fentanyl intake and does not enhance the reinforcing effects of fentanyl. Meanwhile, studies of brain oxygenation have found that naloxone alone is sufficient to reverse the initial hypoxic effects of opioid-xylazine mixtures, but that only naloxone combined with an α2AR antagonist reversed the more prolonged hypoxic effects of fentanyl-xylazine mixtures (Choi et al. 2024). Xylazine blocked the compensatory hyperoxic phase that occurs following the initial opioid-induced hypoxia (Choi et al. 2023), which would explain how xylazine exacerbates the lethal effects of fentanyl. These findings suggest that naloxone and supplemental oxygen may be sufficient to rescue individuals from fentanyl-xylazine overdose, contrary to the widespread notion that xylazine leads to naloxone-resistant overdose, but further research is warranted to better understand the acute effects of fentanyl-xylazine mixtures.

Since our overall understanding of opioid-xylazine interactions is entirely predicated on recent preliminary findings, which have only begun to elucidate these drug interactions at select doses under select experimental conditions, this study sought to broaden the scope of behavioral paradigms being used to examine these interactions. Here we report the effects of opioid-xylazine mixtures using a variety of physiological and behavioral endpoints, including locomotor activity, rectal temperature, subjective effects, and respiration.

## Methods

### Subjects

Adult male Sprague-Dawley rats (Envigo, Indianapolis, IN) were used for these studies. Male rats weighed between 200-220g on arrival. Rats were individually housed in clear open-top cages with wire lids holding chow (Teklad Extruded Global Rodent Diet, Inotiv, Indianapolis, IN) and aspen bedding (Teklad 7093, Inotiv, Indianapolis, IN), maintained on a 12/12-hour light/dark cycle, had access to enrichment (wood gnawing block and rat tunnel), and to food and water *ad libitum* except while behavioral tests were conducted. All behavioral experiments were conducted during the light period between 9:00am and 3:30pm and were conducted by the same female experimenter. Prior to beginning any experimental procedures, each rat was handled by the experimenter for approximately 5 minutes per day for four consecutive days. For all handling and behavioral tests, rats were picked up and carried by hand, and not by the tail. All experiments were performed according to protocols approved by the Institutional Animal Care and Use Committee, University at Buffalo, the State University of New York, and with the *2011 Guide for the Care and Use of Laboratory Animals* (Institute of Laboratory Animal Resources on Life Sciences, National Research Council, National Academy of Sciences, Washington DC).

### Drugs

Fentanyl and morphine sulfate were provided by the Research Technology Branch, National Institute on Drug Abuse, National Institutes of Health (Rockville, MD). Naloxone hydrochloride dihydrate was purchased from Sigma-Aldrich (St. Louis, MO). Xylazine hydrochloride was purchased from MP Biomedicals, LLC (Solon, OH). Atipamezole hydrochloride (Revertidine) was purchased from Modern Veterinary Therapeutics, LLC (Sunrise, FL) as a 5 mg/mL injectable solution. Fentanyl, morphine, naloxone, and xylazine were dissolved in 0.9 percent saline from Dechra Veterinary Products (Overland Park, KS). Atipamezole was diluted using 0.9 percent saline and was injected subcutaneously in a volume of 1 mg/kg in body temperature experiments but was injected intraperitoneally in respiratory experiments where some animals were concurrently administered atipamezole and naloxone. All other drugs were administered intraperitoneally in a volume of 1-2 ml/kg for all experiments.

### Open field test

Locomotion was recorded using an infrared motion-sensor system (AccuScan Instruments, Inc., Columbus, OH) surrounding Plexiglas chambers (40 x 40 x 30 cm). Versa Max software (Omnitech Electronics, Inc., Columbus, OH) was used to quantify the distance each animal traveled throughout the testing period. Male rats (*n* = 78) were habituated to the testing room for 20 minutes (min), injected with a dose of xylazine (0.56, 1, 1.78 or 3.2 mg/kg; *n* = 16 for each treatment dose) or vehicle (*n* = 14), and were immediately placed into the open field chamber for 1 hour. The total distance traveled by each rat over the first 30 min was used for analysis. The final group sizes were not predetermined and are the result of several groups of 8-12 rats undergoing locomotor experiments on separate dates, but all locomotor experiments were conducted during the same time of day (between 10:00am and 1:00pm) and when the rats were approximately the same age (i.e. 6-10 days after arrival). Group sizes were larger than the 4-6 animals per treatment condition that were initially planned on account of variability in sensitivity to xylazine observed between animals. All animals were repurposed for other behavioral experiments, some of which are described in the following sections, but were given a minimum of 5 days prior to beginning other pharmacological experiments to allow time for drug clearance.

### Measurement of body temperature

Body temperature measurements were collected in a quiet testing room with the same environmental conditions as the housing room between 9:00am and 2:00pm. Two groups of rats (*n* = 6 per group) were handled by the experimenter for at least 4 days, and 2 saline sessions were conducted prior to testing to allow for rats to adjust to the experimental procedure. On testing days, after 20-30 min of habituation to the testing room, a portable temperature monitor (PTM1) and rectal probe (RET-2) from Physitemp Instruments (Clifton, NJ) were used to measure each subject’s body temperature before and after receiving an intraperitoneal injection with a drug treatment. In the reversal experiment, atipamezole was administered subcutaneously 90 min after initial treatment with xylazine. Measurements were recorded every 15 min for a total duration of 3 hours. Body temperature experiments were conducted such that 3 or more days elapsed between behavioral tests, to allow time for drug clearance and reduce the likelihood of drug accumulation in subjects.

### Drug discrimination

Operant studies were conducted using two-lever Habitest chambers in sound-attenuating enclosures (Coulbourn Instruments LLC, Allentown, PA), which were connected to a LabLinc interface and a computer running Graphic State software (version 3.03). Rats initially underwent daily shaping sessions made up of two cycles, where each cycle consisted of a 10 min timeout period, followed by a 5 min response period, during which a house light and cue lights above the levers were illuminated and up to 5 food reinforcers could be earned. Rats began at a fixed ratio (FR) 1, where 1 lever press would earn 1 food pellet (45 mg, Bio-Serv, Flemington, NJ) and progressed up to FR10 over the course of about a week. The lever that was reinforced was altered between left and right each day, until rats consistently responded at FR10 on both the left and right lever. Training sessions consisted of the same 2 cycles at FR10, except the correct lever was determined by the injection that was administered intraperitoneally immediately prior to beginning the session (i.e., right lever, saline; left lever, training drug). Training sessions were conducted 5-7 days a week using a double alternation schedule (i.e., saline, saline, drug, drug) until rats met the following criteria for 5 consecutive or 6 out of 7 consecutive sessions: 1) less than 10 responses were made on the incorrect lever prior to earning the first food reward, and 2) at least 80% of total responses made during the active periods were on the correct lever. Thereafter, test sessions were conducted using the same procedure, except 10 responses on either lever resulted in delivery of a food pellet. Between test sessions, rats had to meet the same criteria for another 2 consecutive sessions (a saline session and a training drug session). Animals were placed into the operant chamber immediately after being injected with drug or saline and therefore began responding 10 min after drug administration, except in pretreatment experiments where animals were injected with drug and returned to their home cages in the testing room until the pretreatment time (*t* - 10 min to account for the 10 min timeout period) had elapsed. Tests were repeated if an animal’s results were inconsistent with other doses tested (for example, if an animal responded 100% on the fentanyl lever at 2 mg/kg morphine but at 50% for a dose of 4 mg/kg morphine) or if an animal did little to no responding at a dose of drug but later responded at higher doses in subsequent tests (i.e. the subject elected not to participate but was not necessarily sedated). If the animal did not earn a reward during a training session, then data from a subsequent test where the animal did respond was used in data analysis. Otherwise, data collected in separate test sessions was averaged. Calculations of sessions to criteria excluded days that animals were too sedated or elected not to engage in the task. Data from training sessions can be found in **Supplemental Fig. 1**.

A group of 12 rats was initially trained using a dose of 0.04 mg/kg fentanyl, but 5 animals that exhibited little to no lever responding at a dose of 0.04 mg/kg fentanyl (owing to rate-suppressant effects) were instead moved to a training dose of 0.02 mg/kg fentanyl. 2 animals that met or were close to meeting criteria at the 0.02 mg/kg training dose were removed from the study due to health complications. 2 animals that were trained to discriminate 0.04 mg/kg fentanyl from saline were later retrained to discriminate 0.02 mg/kg fentanyl for conducting experiments with added xylazine. Thus the 0.04 mg/kg group consisted of *n* = 7, and the 0.02 mg/kg group consisted of *n* = 5.

A group of 8 rats was initially trained using a dose of 1.5 mg/kg xylazine, but 3 animals were later moved to a training dose of 1 mg/kg xylazine due to rate-suppressant effects. 1 animal was removed from the study because food restriction was insufficient motivation to facilitate consistent lever pressing at FR10 and his sporadic participation made it difficult to assess his learning. Therefore the 1.5 mg/kg group consisted of *n* = 4. 1 rat met criteria at the 1 mg/kg training dose but was not included in this study.

### Whole body plethysmography

Respiratory behavior was measured using two rat-sized whole body plethysmography chambers (Buxco Small Animal WBP, Data Sciences International, St. Paul, MN) connected to a computer (Intel Core i5, Dell) with FinePointe Software. Both chambers were individually calibrated on each test day prior to beginning a recording. Each rat was placed in the plethysmography chamber with the equipment running for 30 min a day for 2 days to habituate to the equipment. Then each rat underwent the same procedure but received an intraperitoneal injection of saline after 30 min elapsed and was returned to the chamber for an additional 30 min for 3 more days to habituate the animals to the experimental protocol. Thereafter, rats were administered with a pharmacological treatment once a week using a randomized Latin square design, where treatment groups consisted of 4-6 individuals from a pool of rats. Each experiment was constrained to a 2–3-week period, since tidal volume changes occur with aging and increasing body weight.

A pool of 12 rats was used to collect data for xylazine alone and fentanyl-xylazine co-administration experiments, morphine-xylazine co-administration experiments, and naloxone reversal experiments. A separate pool of 12 rats – initially used for body temperature experiments – was used for atipamezole and naloxone reversal experiments. There was a drug free period of at least 2 weeks between the time body temperature experiments took place and respiratory depression reversal tests began. An additional pool of 12 rats were used to conduct atipamezole blockade/reversal experiments.

Recordings for xylazine alone, fentanyl-xylazine co-administration, morphine-xylazine co-administration consisted of a 30 min baseline period in the chamber, then animals were briefly removed from the chamber and injected intraperitoneally before being returned to the chamber for an additional 1-2 hour recording period. In reversal experiments, animals were injected prior to beginning the first 30 min recording period, and were removed and injected again before being returned to the chamber for an additional 1-2 hour recording period.

### Data analyses

Locomotion and body temperature data were analyzed in GraphPad Prism (GraphPad Software, La Jolla, CA) using an ordinary one-way analysis of variance (ANOVA) and a two-way repeated measures analysis of variance (rmANOVA), respectively. These were each followed by a post hoc Bonferroni’s multiple comparisons test. The threshold for significance was set at P≤0.05.

Drug discrimination data are shown as the average percentage of responses made on the drug-lever and the average response rate during the active period using both cycles within the session. This data was plotted for each individual subject, in addition to being expressed as the mean ± SEM for the group of rats. Percent of responding on the drug-lever at a particular dose was only shown if a rat earned at least 5 rewards during the test session, and data for the group was only shown if at least half of the group earned a minimum of 5 rewards. The response rates are shown irrespective of the number of rewards earned. For 0.04 mg/kg fentanyl and 1.5 mg/kg xylazine drug discrimination experiments, the median effective dose (ED_50_) values and 95% confidence intervals were calculated using linear regression. The ascending limb of an inverted U-shaped dose response curve was used to calculate the ED_50_ where applicable. Comparisons between drug treatment and control (i.e. saline, or 0.02 mg/kg fentanyl) groups were made with multiple paired t-tests using the Bonferroni-Dunn method. Drug-lever responding and rate of responding data from the saline session prior to each test were used for statistical analysis.

Respiratory measures for frequency, minute volume, and tidal volume for each individual animal were averaged for each min of the recording within FinePointe. These values were input into GraphPad Prism to be visualized and to conduct statistical analysis. In time course graphs, the control data is expressed as mean ± 95 percent confidence interval and the data from each treatment dose are mean ± SEM. Area under the curve for specific bins of time were calculated in GraphPad Prism and the resulting bar graphs show the mean ± 95 percent confidence interval for each treatment group, differences between treatment groups were considered statistically significant when the 95 percent confidence intervals had no overlap.

## Results

Intraperitoneal xylazine treatment significantly decreased locomotor activity in an open field at doses of 0.56, 1, 1.78, and 3.2 mg/kg (one-way ANOVA: F_(4, 73)_ = 7.111, P<0.0001) (**Fig. 1**). Rather than incremental dose-dependent decreases the sedative effects of xylazine exhibited a plateau-like effect, likely due to the considerable variability between subjects. The observation that some animals were unaffected by high doses of xylazine, while others were sedated at low doses led us to parse our other behavioral data at the group and individual levels.

**Fig. 1.**
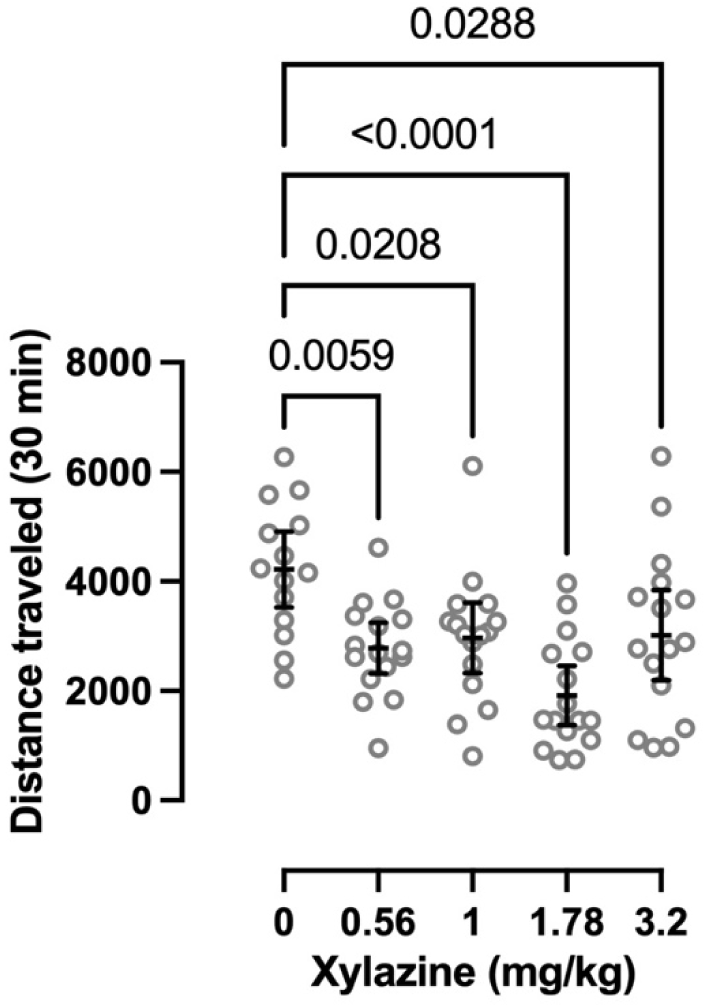
Effects of acute xylazine treatment on locomotor activity. Each circle indicates the cumulative distance an individual animal travelled in an open field during the 30 min period immediately after i.p. administration. Overlayed bars indicate mean ± 95% confidence interval. P-values were determined using an ordinary one-way ANOVA followed by Bonferroni’s multiple comparisons test. Saline/vehicle group consisted of *n* = 14; xylazine treatment groups were *n* = 16.

Clonidine, an α2AR agonist, has previously been shown to dose-dependently decrease rectal temperature in rats, therefore we evaluated the effects of xylazine on body temperature. Xylazine decreased body temperature at doses of 5.6 and 10 mg/kg, but its effects were more variable and less potent than clonidine (**Fig. 2a**). When xylazine was administered with morphine, there were no significant differences from xylazine alone (**Fig. 2b**), but there was a main effect of time (two-way RM ANOVA: F_(11, 165)_ = 22.18, P<0.0001) and subject (F_(15, 165)_ = 44.16, P<0.0001). In contrast, when clonidine was administered with morphine, there was a significant interaction between time and treatment (two-way RM ANOVA: F_(22, 165)_ = 6.179, P<0.0001), along with a main effect of time (F_(11, 165)_ = 248.2, P<0.0001) and subject (F_(15, 165)_ = 15.87, P<0.0001). Post hoc analysis found that the addition of 10 mg/kg morphine significantly increased body temperature between 30 and 90 min after drug administration, and the addition of 1 mg/kg morphine significantly decreased body temperature at 150 and 180 min after drug administration. The hypothermic effects of 10 mg/kg xylazine were reversed by the α2AR antagonist atipamezole (**Fig. 2c**), suggesting this effect is mediated at least in part by α2ARs similar to clonidine. There was a main effect of time (two-way RM ANOVA: F_(11, 220)_ = 21.55, P<0.0001), treatment (F_(3, 20)_ = 15.74, P<0.0001), and subject (F_(20, 220)_ = 25.21, P<0.0001), in addition to a significant interaction between time and treatment (F_(33, 220)_ = 8.550, P<0.0001). Post hoc analysis found that 0.1 mg/kg atipamezole significantly increased body temperature between 30 and 75 min after drug administration, whereas 1 mg/kg atipamezole significantly increased body temperature between 15 and 90 min after drug administration. The hypothermic effects of 10 mg/kg xylazine varied considerably between individuals – the body temperature of one animal was unaffected, one had more pronounced hypothermic effects, and the body temperature of the remaining four animals dropped between 2-4 degrees (**Fig. 2d**). The addition of 1 mg/kg morphine appeared to have either no effect or modestly decreased body temperature, while the addition of 10 mg/kg morphine attenuated the hypothermic effects of xylazine in some subjects.

**Fig. 2.**
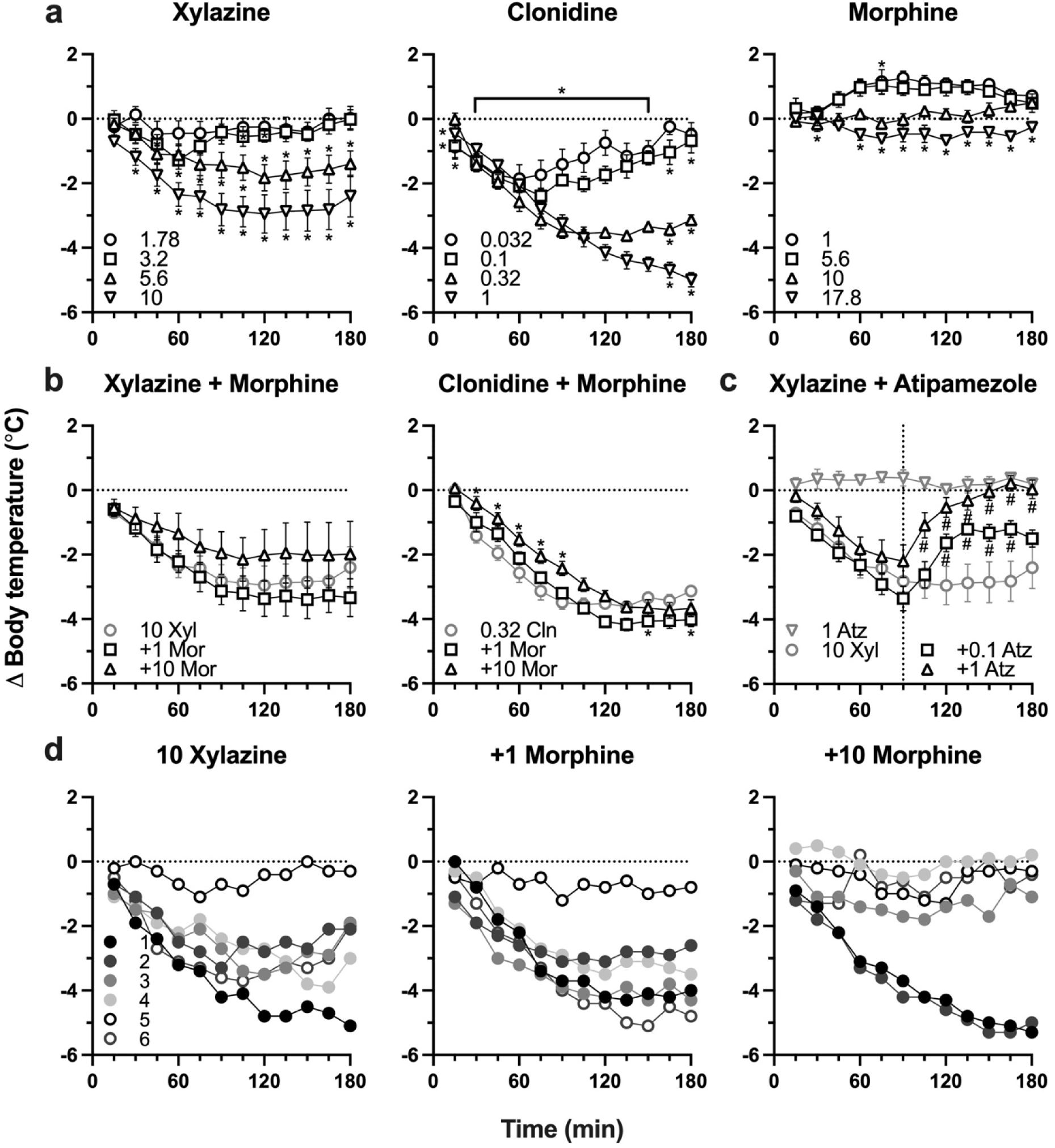
Effects of α_2_ adrenergic agonists and morphine on rectal temperature. **a**) Body temperature changes induced by treatment with xylazine, clonidine, and morphine alone. **b**) Changes in body temperature resulting from treatment with both an α_2_ adrenergic agonist and morphine. **c**) Reversal of xylazine effects on body temperature by α_2_ adrenergic antagonist atipamezole. Data were analyzed using a two-way repeated measures ANOVA followed by Bonferroni’s multiple comparisons test, and symbols (* or #) indicate time points that were significantly different from xylazine or clonidine alone. Each treatment group consisted of *n* = 6. **d**) Effects of xylazine and xylazine + morphine on individual animals. Legends indicate dose in mg/kg (**a**, **b**, **c**) or subject (**d**).

Seven rats acquired discrimination between 0.04 mg/kg fentanyl and saline in an average (± SEM) of 70.6 (± 7.4) training sessions. Fentanyl dose-dependently increased responding on the fentanyl-associated lever up to an average maximum of 99.0% at a dose of 0.08 mg/kg (**Fig. 3a**, open circles). Morphine similarly resulted in a dose-dependent increase in fentanyl-associated lever responding up to an average maximum of 96.6% at a dose of 8 mg/kg (**Fig. 3a**, open squares). Xylazine occasioned responding on the fentanyl-associated lever up to an average maximum of 37.5% at a dose of 1 mg/kg (**Fig. 3a**, open triangles), with individual animals responding between 0% and 97.2% on the fentanyl-associated lever. The ED_50_ values (95% confidence intervals) for the discriminative stimulus effects of fentanyl and morphine were 0.0139 mg/kg (0.0088-0.0190 mg/kg) and 3.369 mg/kg (2.159-4.579 mg/kg), respectively.

**Fig. 3.**
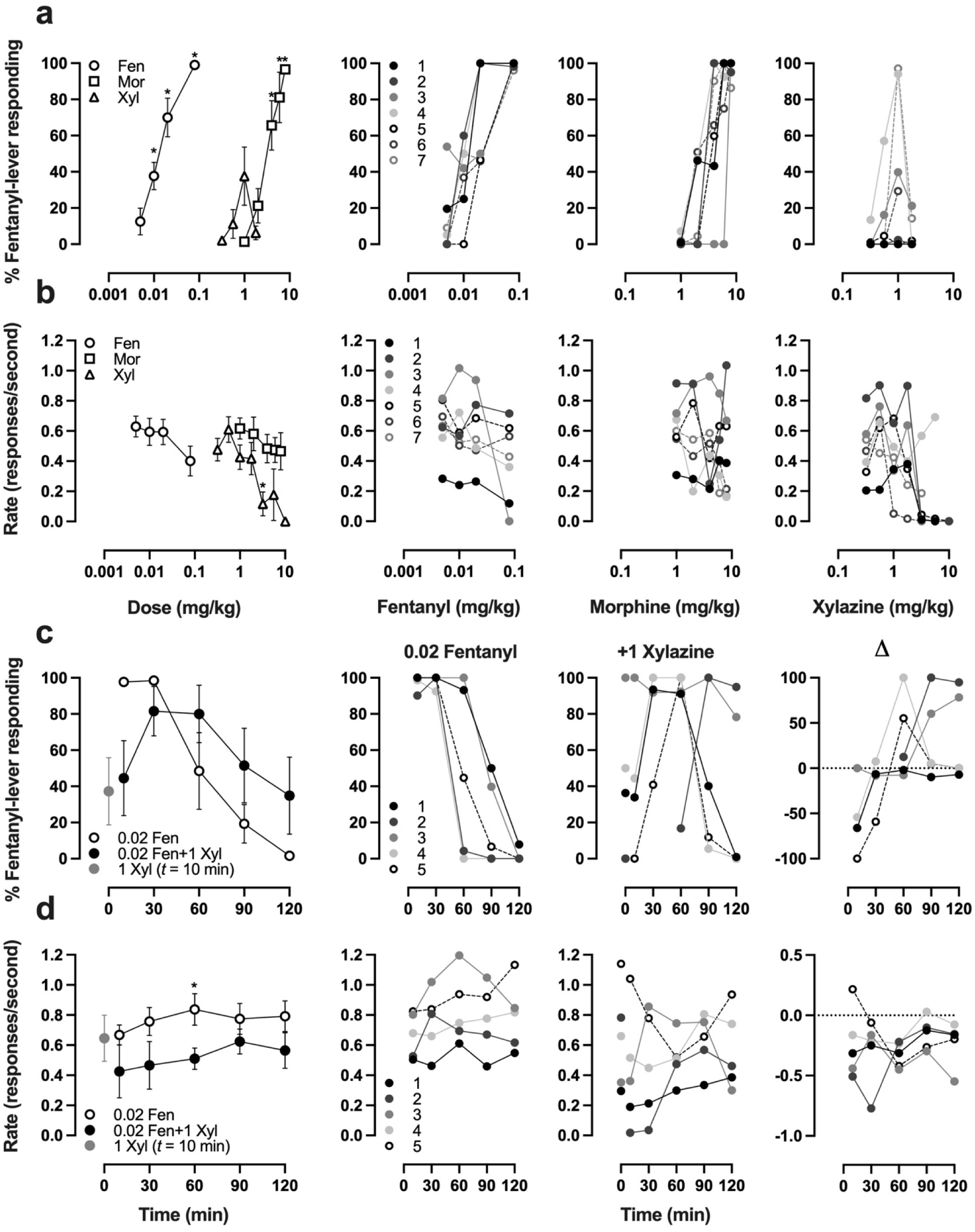
Effects of xylazine on animals trained to discriminate fentanyl from saline. Discriminative stimulus effects (**a**) and rate suppressant effects (**b**) of fentanyl, morphine, and xylazine on rats trained to discriminate 0.04 mg/kg fentanyl from saline. *Left:* Mean ± SEM of 7 animals where asterisks indicate significant difference from saline sessions. *Right:* Results of individual animals at different doses of fentanyl (*left*), morphine (*center*), and xylazine (*right*). Pharmacokinetic effects of fentanyl and fentanyl-xylazine co-treatment on fentanyl discriminative stimulus (**c**) and rate responding (**d**) in rats trained to discriminate 0.02 mg/kg fentanyl from saline. *Left:* Mean ± SEM of 5 animals where asterisks indicate a significant difference from 0.02 mg/kg fentanyl. *Right:* Results of individual animals when treated with fentanyl (*left*), xylazine or fentanyl and xylazine (*center*), and the difference between fentanyl-xylazine and fentanyl (*right*).

Three of five rats acquired discrimination between 0.02 mg/kg fentanyl and saline in an average (± SEM) of 123.3 (± 4.5) training sessions, and the two rats that were retrained from 0.04 mg/kg fentanyl reacquired discrimination at a lower dose after an additional 18.0 (± 7.0) training sessions. As fentanyl pretreatment times increased, the average fentanyl-associated lever responding decreased, and two hours after receiving the training dose the average maximum responding on the fentanyl-associated lever was 1.6% (**Fig. 3c**, open circles). The addition of 1 mg/kg xylazine produced a nonsignificant increase in fentanyl-lever responding (one-tailed paired t-test: P = 0.3868, t = 0.3078, df = 4) and a significant decrease in response rate (P = 0.0006, t = 8.268, df = 4), with considerable variability between animals (**Fig. 3c** and **3d**). When probed with 1 mg/kg xylazine alone, xylazine occasioned responding on the fentanyl-associated lever up to an average maximum of 37.3%, with individuals responding between 0% and 100% on the fentanyl-lever.

Four rats acquired discrimination between 1.5 mg/kg xylazine and saline in an average (± SEM) of 96.8 (± 22.0) training sessions. Responding on the xylazine-associated lever peaked at an average maximum of 86.9% at the training dose of 1.5 mg/kg (**Fig. 4a**, open circles), but decreased at doses above the training dose. Clonidine occasioned responding on the xylazine-associated lever up to an average maximum of 96.9 and 98.2% at doses of 0.032 and 0.056 mg/kg, respectively (**Fig. 4a**, open squares). Fentanyl occasioned responding on the xylazine-associated lever up to a maximum of 22.6% at a dose of 0.02 mg/kg (**Fig. 4a**, open triangles), with individual animals responding between 0% and 48.1% on the xylazine-lever. The ED_50_ values (95% confidence intervals) for the discriminative stimulus effects of xylazine and clonidine were 0.894 mg/kg (0.449-1.338 mg/kg) and 0.0153 mg/kg (0.0058-0.0248 mg/kg), respectively.

**Fig. 4.**
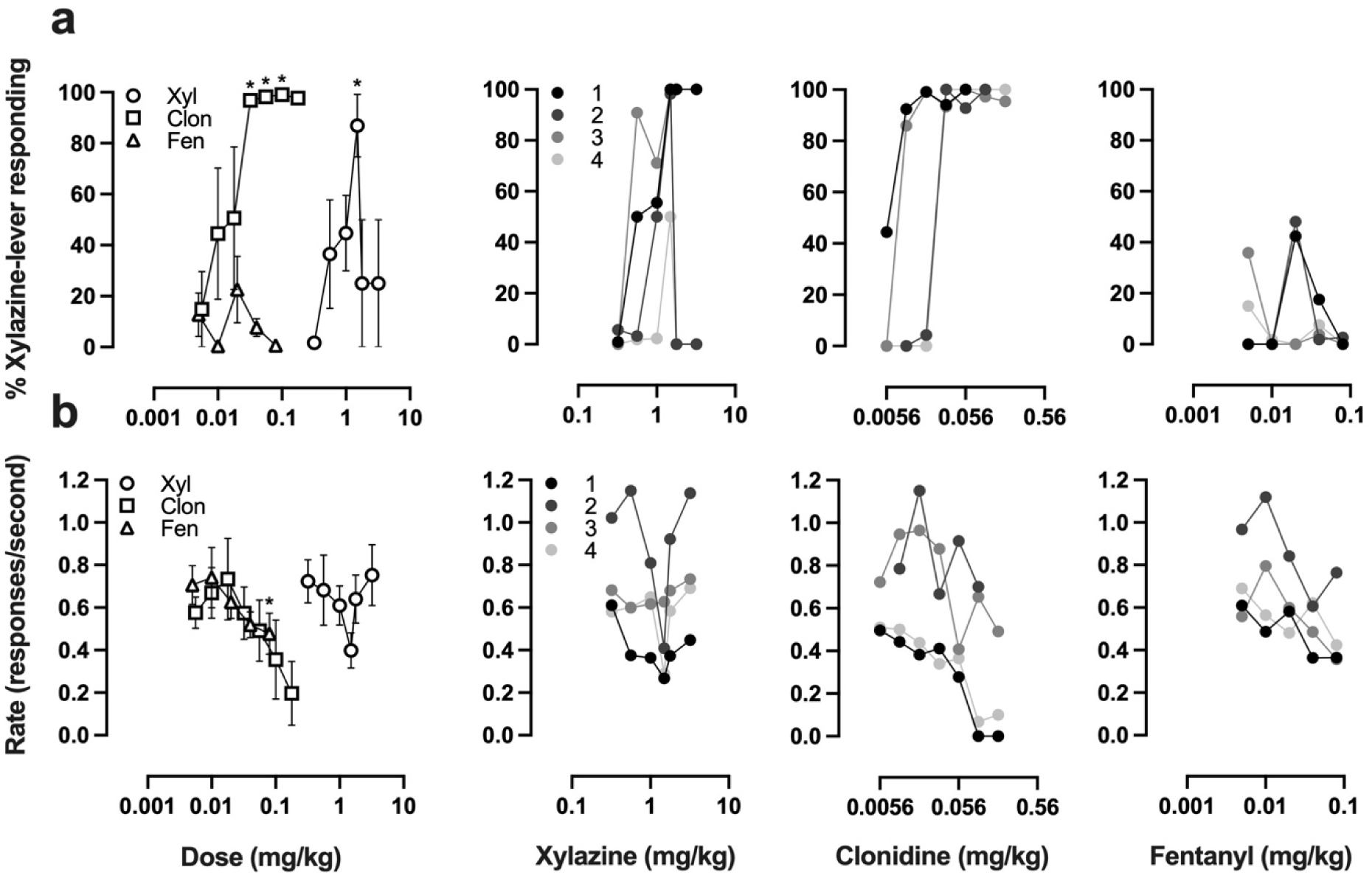
Effects of fentanyl on animals trained to discriminate xylazine from saline. Discriminative stimulus effects (**a**) and rate suppressant effects (**b**) of xylazine, clonidine, and fentanyl on rats trained to discriminate 1.5 mg/kg xylazine from saline. *Left:* Mean ± SEM of 4 animals where asterisks indicate significant difference from saline sessions. *Right:* Results of individual animals at different doses of xylazine (*left*), clonidine (*center*), and fentanyl (*right*).

Although preclinical studies have demonstrated that xylazine potentiates the lethality of fentanyl and potentiates the hypoxic effects of opioids in specific brain regions following intravenous administration, knowledge of the extent to which xylazine exacerbates opioid-induced respiratory depression is limited. To evaluate the effects of xylazine alone and when co-administered with opioids, we utilized whole body plethysmography to measure changes in respiratory frequency, minute volume, and tidal volume in unrestrained rats. Intraperitoneal injection of xylazine at doses of 1 mg/kg and 1.78 mg/kg transiently decreased minute volume 0-10 min after administration, and 1.78 mg/kg xylazine increased tidal volume between 20-60 min post-injection (**Fig. 5a**). 0.56 mg/kg xylazine increased respiratory frequency and minute volume 20-60 min after injection. When 10 mg/kg morphine and 1 mg/kg xylazine were co-administered there was a decrease in respiratory frequency 0-10 min after injection, a decrease in tidal volume between 10-40 min post-injection, and a decrease in minute volume 40-60 min after administration (**Fig. 5b**). Interestingly, the addition of 1.78 mg/kg xylazine had no significant effects during the period of observation, perhaps due to compensatory effects or due to the randomized group of subjects being less sensitive to the respiratory effects of xylazine than the group that received 1 mg/kg. When 0.1 mg/kg fentanyl and 1.78 mg/kg xylazine were co-administered respiratory frequency decreased 10-60 min post-injection, and tidal volume increased 20-60 min after administration (**Fig. 5c**). When fentanyl was co-administered with 0.56 mg/kg xylazine respiratory frequency and minute volume increased between 20-40 min post-injection. The finding that the addition of 1.78 mg/kg xylazine to 0.1 mg/kg fentanyl led to a prolonged decrease in respiratory frequency suggests xylazine may exacerbate fentanyl-induced respiratory depression, however there was a compensatory increase in tidal volume, which prevented any notable changes in minute volume. Furthermore, since 0.56 mg/kg xylazine increased respiratory frequency and minute volume when administered both alone and alongside 0.1 mg/kg fentanyl, it may be that whether xylazine exacerbates opioid-induced respiratory depression is dependent on the dose of xylazine or on the individual.

**Fig. 5.**
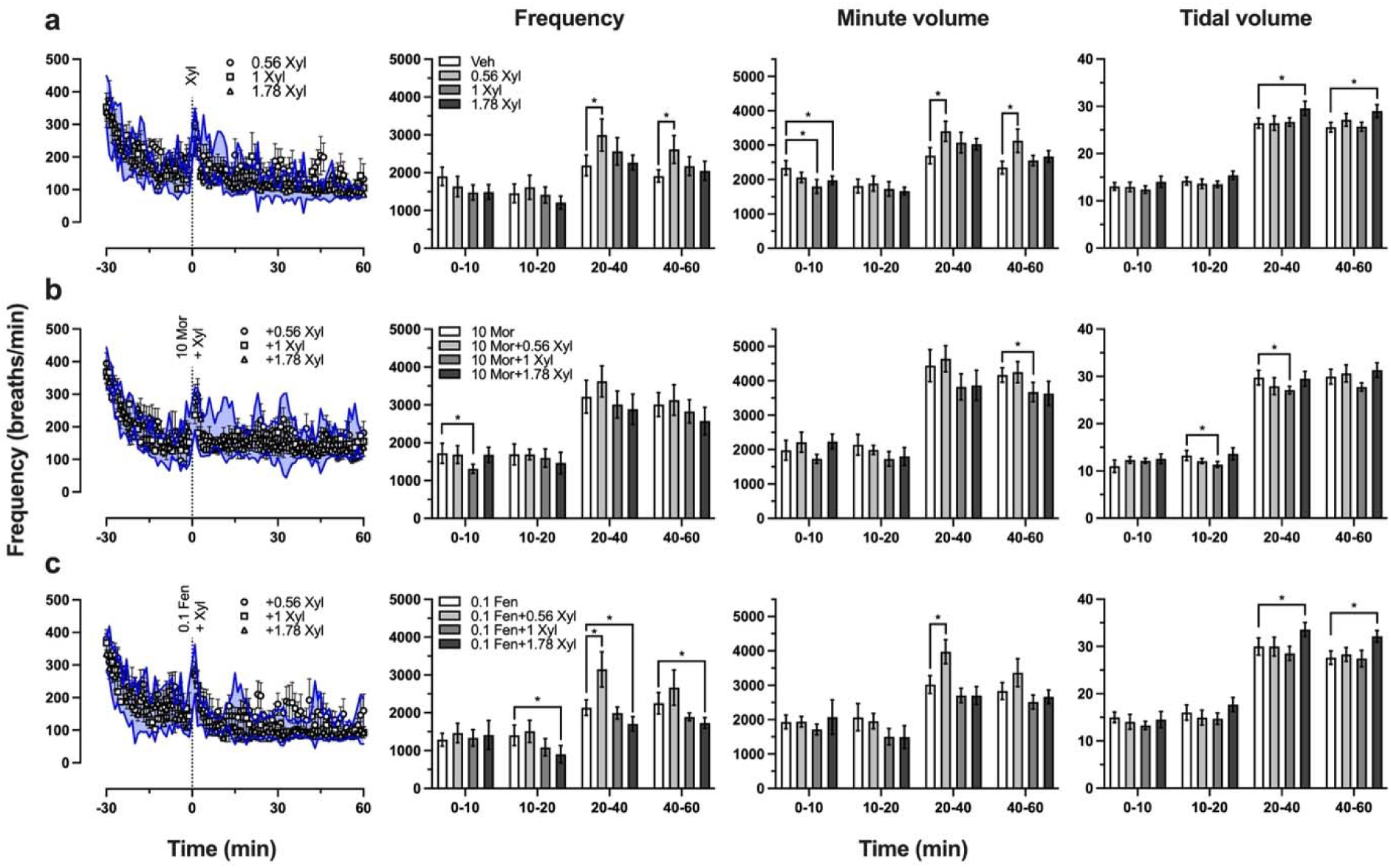
Effects of xylazine, fentanyl-xylazine co-administration, and morphine-xylazine co-administration on respiration. *Left:* Respiratory frequency (breaths per minute) before and after drug treatment, where the blue shaded region is the 95% confidence interval of saline-treated (**a**), 10 mg/kg morphine-treated (**b**), or 0.1 mg/kg fentanyl-treated (**c**) animals. *Right:* Bar graphs show area under the curve calculated for frequency (*left*), minute volume (*center*), and tidal volume (*right*) from specific bins of time, and are depicted as mean ± 95% confidence interval. Asterisks indicate instances where there is no overlap in 95% confidence intervals. Control groups consisted of *n* = 6, all other groups consisted of *n* = 4. Legends indicate dose of drug(s) tested in mg/kg.

Since the idea that xylazine leads to naloxone-resistant overdose is widely circulated despite a lack of preclinical evidence, we again utilized whole body plethysmography to examine the efficacy of naloxone, atipamezole, and concurrent naloxone and atipamezole at reversing the respiratory effects of opioids and xylazine. When 10 mg/kg morphine and 1.78 mg/kg xylazine were administered 30 min prior to 1 mg/kg naloxone, the only significant differences between this group and the group that received only 10 mg/kg morphine and naloxone 30 min thereafter was seen in tidal volume between 10-40 min post-naloxone (40-70 min after morphine or morphine-xylazine administration) (**Fig. 6a**). Although 10 mg/kg morphine and 1.78 mg/kg xylazine had no significant effects in the prior group (**Fig. 5b**), the control group that received 10 mg/kg morphine and 1.78 mg/kg xylazine and vehicle 30 min later exhibited significantly decreased respiratory frequency (30-40 and 50-70 min after morphine-xylazine) and minute volume (30-90 min after morphine-xylazine), suggesting the effect of xylazine on opioid-induced respiratory depression may depend on the individual. When animals treated with 10 mg/kg morphine and 1.78 mg/kg xylazine were administered 1 mg/kg naloxone, 1 mg/kg atipamezole, or 1 mg/kg naloxone and 1 mg/kg atipamezole, we found that atipamezole alone was as effective at reversal as naloxone-atipamezole combination (**Fig. 6b**). Naloxone alone increased respiratory frequency and minute volume between 0-10 min after injection, and increased tidal volume between 20-60 min post-injection, which suggests naloxone is an effective reversal agent, but the effect is transient. In contrast, atipamezole alone and co-administered with naloxone led to a prolonged increase in respiratory frequency and minute volume. When the respiratory frequency data from **Fig. 6b** is shown by individual animal, we observed that a few individuals were unaffected by co-administration of morphine and xylazine (**Fig. 6c**), supporting the idea that xylazine exacerbates opioid-induced respiratory depression in some individuals but not others. Since atipamezole is an α2AR antagonist and appeared to effectively antagonize morphine-xylazine induced respiratory depression effectively by itself (**Fig. 6b** and **6c**), we then examined whether atipamezole was able to block or reverse the respiratory effects of morphine and fentanyl without the addition of xylazine. When 0.1 mg/kg fentanyl and 1 mg/kg atipamezole were administered at the same time, atipamezole appeared to block the fentanyl-induced depression of respiratory frequency, whereas atipamezole treatment 30 min after fentanyl injection resulted in a transient (∼15 min) reversal effect (**Fig. 6d**). Atipamezole did not have any observable effect on respiration frequency when administered alongside or 30 min after 10 mg/kg morphine (**Fig. 6e**), though the respiratory effects of 10 mg/kg morphine were less pronounced and more variable than 0.1 mg/kg fentanyl and therefore may have precluded any observable effect.

**Fig. 6.**
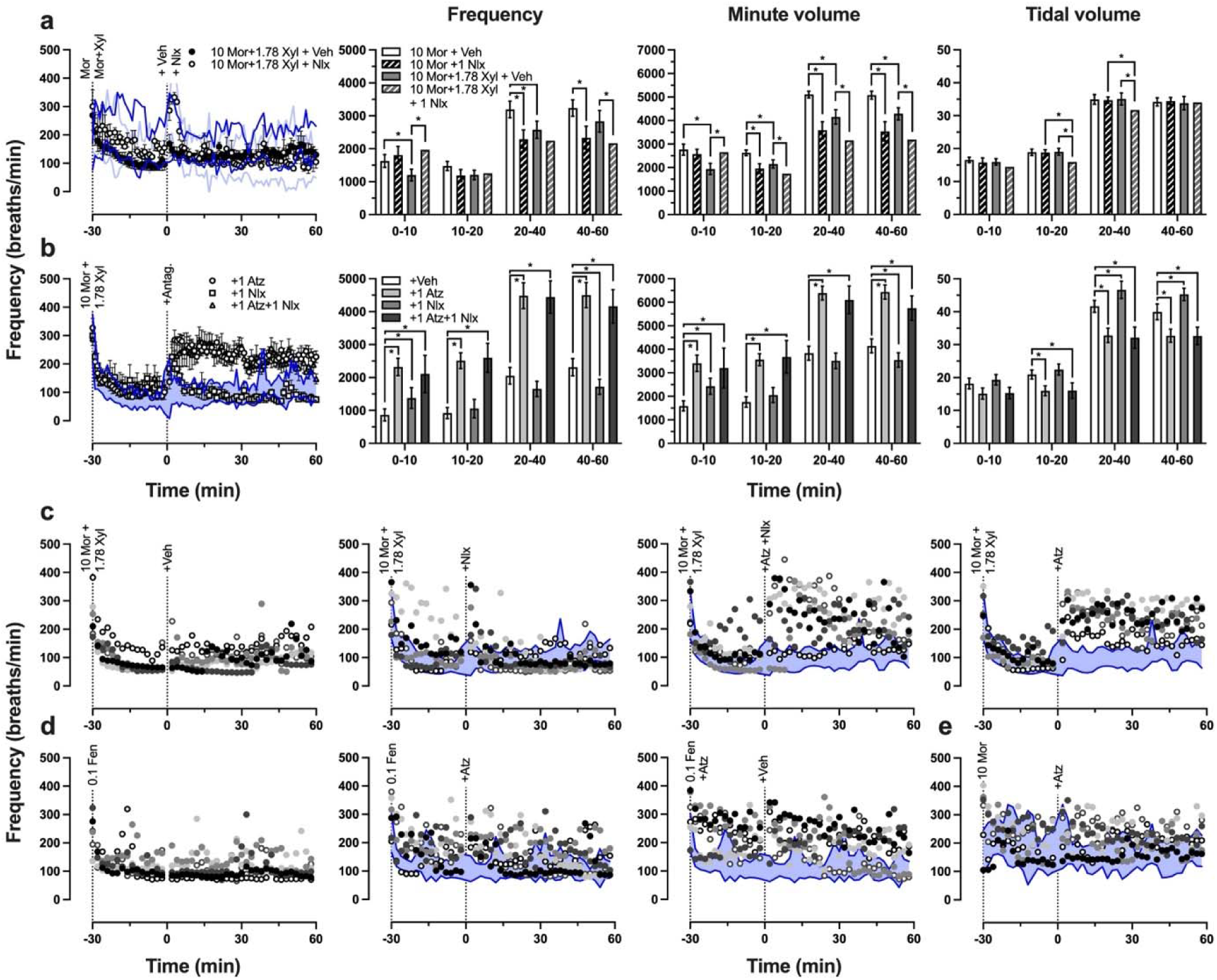
Effects of naloxone, naloxone-atipamezole co-administration, and atipamezole on opioid-induced respiratory depression. **a**) *Left:* Respiratory frequency (breaths per minute) following morphine or morphine-xylazine treatment and administration of naloxone or vehicle 30 min thereafter, where the dark blue lines indicate the 95% confidence interval of 10 mg/kg morphine followed by vehicle treatment and the light blue lines indicate the 95% confidence interval of 10 mg/kg morphine followed by 1 mg/kg naloxone treatment. *Right:* Area under curve calculated for frequency (*left*), minute volume (*center*), and tidal volume (*right*) for specific intervals of time after the antagonist or vehicle control was administered. Each group consisted of *n* = 4. **b**) *Left:* Frequency of respiration following treatment with 10 mg/kg morphine and 1.78 mg/kg xylazine and administration of an antagonist or vehicle, where blue shaded region is 95% confidence interval of vehicle-treated animals. *Right:* Quantification of area under curve following treatment with antagonist or vehicle control, asterisks indicate instances where there is no overlap in 95% confidence intervals. Each group consisted of *n* = 6. **c** and **d**) Respiratory frequency of individual animals over time, where leftmost graph shows the animals that were used for the blue shaded 95% confidence intervals in rightward panels. **e**) Respiratory effects of 10 mg/kg morphine followed by 1 mg/kg atipamezole, where control group is *n* = 4 and treatment group is *n* = 6. Note that reuse of the same symbols in separate panels does not represent the same animal, as these groups were randomized via Latin square design.

## Discussion

The primary finding of this study was not that there were marked individual differences in any singular behavioral readout, but rather that dramatic interanimal variability repeatedly emerged in a variety of behavioral assessments despite experimental groups consisting of a relatively small number of male Sprague Dawley rats. This is perhaps most appreciable in the reported locomotor effects of xylazine (**Fig. 1**) where the variability between animals precludes determination of an ED_50_ for the locomotor suppressant effect of xylazine. Similar results have been observed in C57BL/6J mice, where 3 mg/kg xylazine (i.p.) significantly decreased locomotor activity, but the locomotor effects of 0.5 and 1 mg/kg xylazine on individual mice were highly variable (Bedard et al. 2024). Significant depression of locomotor activity has been reported in male mice treated with the α2AR agonist clonidine (0.05-0.5 mg/kg, i.p.) (Capasso et al. 1996; Capasso and Loizzo 2001), whereas 1 mg/kg clonidine (i.p.) did not significantly reduce locomotor activity in male Wistar rats (Ozcetin et al. 2016). Interestingly, while clonidine (1-10 mg/kg, i.p.) significantly suppressed locomotor activity in shrews (Sun and Darmani 2024), the results were highly variable and thus exhibited a plateau-like effect similar to what we observed with xylazine. Altogether, these findings highlight the disparate effects of α2AR agonists on locomotor activity across individuals and species.

α2AR agonists such as clonidine and dexmedetomidine have established hypothermic effects, which are mediated by α2AR and imidazoline receptors (Bhalla et al. 2011; Lahdesmaki et al. 2003). Our finding that xylazine-induced hypothermia was partially attenuated by a selective α2AR antagonist is consistent with these prior findings. Furthermore, the observation that the decrease in body temperature elicited by clonidine plus morphine was similar to what was observed with clonidine alone recapitulates what was reported by Bhalla et al. in 2011. Notably, one subject was consistently insensitive to the hypothermic effects of xylazine, while another subject was consistently more sensitive to the hypothermic effects of xylazine than the rest of the cohort. In contrast, the hypothermic effects of clonidine and clonidine plus morphine were remarkably similar across all subjects.

The discriminative stimulus effects of xylazine and fentanyl plus xylazine are largely uncharacterized. A recent study describes training male and female Long Evans rats to discriminate 0.032 mg/kg fentanyl from saline and found that xylazine did not substitute for fentanyl – although females exhibited some responding on the fentanyl-lever after treatment with 1 mg/kg xylazine – and reported that xylazine potentiated the subjective effects of low doses of fentanyl (Bender et al. 2025). In male Sprague Dawley rats trained to discriminate 0.04 mg/kg fentanyl from saline, we found that 1 mg/kg xylazine partially or even fully substituted for fentanyl in some subjects (**Fig. 3a**). These disparate results may be due to differences in strain, in training/testing sessions (2 cycles of 10 min timeout followed by a 5 min active period vs 10 min timeout and up to 15 min active period), in analysis (average percent responding across 2 cycles vs percent responding prior to first reinforcer), or in individuals that comprise the experimental groups. Though we did not use a 30 min pretreatment time nor administered 0.04 mg/kg fentanyl subcutaneously like past drug discrimination studies, our rats met criteria within 70.6 (± 7.4) sessions, which is in between the 48 and 79 ± 5 STC reported by other groups (De Vry et al. 1984; Zhang et al. 2000). Furthermore, our reported ED50s of 0.0139 (0.0088-0.0190) mg/kg fentanyl and 3.369 (2.159-4.579) mg/kg morphine are consistent with 0.016-0.026 mg/kg fentanyl and 2.98 mg/kg morphine reported by De Vry et al., and 0.021 (0.016, 0.030) mg/kg fentanyl and 3.0 (2.6, 3.4) mg/kg morphine reported by Zhang et al.

Instead of combining low doses of fentanyl with different doses of xylazine to assess the effect on fentanyl-responding like Bender et al., we pretreated animals with a fixed dose of fentanyl (the training dose of 0.02 mg/kg) and a fixed dose of xylazine (1 mg/kg) and initiated the test session at various intervals (0, 20, 50, 80, and 110 min) to evaluate whether the addition of xylazine prolongs the duration of fentanyl’s subjective effects. Overall, we found that 1 mg/kg xylazine initially decreased the subjective effects of fentanyl, but that the addition of xylazine produced a nonsignificant increase in the duration that animals responded on the fentanyl-lever (**Fig. 3c**). The nonsignificant increase was likely due to considerable variability between subjects, where one animal only exhibited a decreased response rate, one animal had no change in fentanyl-lever responding 10 min after receiving both fentanyl and xylazine, and one animal had a more prolonged period of rate depressant effects followed by considerable fentanyl-lever responding at later time points. Altogether these results lend some support to anecdotal evidence from human subjects that xylazine prolongs the subjective effects of fentanyl (Friedman et al. 2022), but also suggest this may not be the case for some individuals.

Conversely, to address whether fentanyl can substitute for xylazine we trained a cohort of male rats to discriminate 1.5 mg/kg xylazine from saline. We chose a training dose of 1.5 mg/kg xylazine as a compromise between rate-suppressant effects and a discriminable dose based on a prior xylazine drug discrimination study (Colpaert and Janssen 1985). Two of four rats treated with 0.02 mg/kg fentanyl responded >40% on the xylazine-lever, but no other dose elicited partial substitution for 1.5 mg/kg xylazine in any of the subjects. Interestingly, xylazine exhibited an inverted U-shaped dose response curve, where animals responded primarily on the saline-lever for doses both above and below the training dose (**Fig. 4a**). Clonidine fully substituted for xylazine in all four subjects, suggesting these animals were able to effectively discriminate the subjective effects of α2AR agonists from saline but that the subjective effects of xylazine at doses above or below the training dose of 1.5 mg/kg xylazine are markedly different. The prior xylazine discrimination study used a 30 min pretreatment time, and consisted of a 15 min active period, but nevertheless reported that variability among animals was large (Colpaert and Janssen 1985), which aligns with our observations. Although animals in the prior study met criteria more quickly (50.4 ± 7.4 vs 96.8 ± 22.0 STC), the reported ED50s for animals discriminating 2.5 mg/kg and 1.25 mg/kg xylazine were 1.03 and 0.70 mg/kg, respectively, which aligns with our reported ED50 of 0.894 mg/kg for rats discriminating 1.5 mg/kg xylazine.

The inverted U-shaped xylazine dose response curve was seen in both fentanyl-discriminating (**Fig. 3a**) and xylazine-discriminating rats (**Fig. 4a**). Inverted U-shaped dose response curves have also been observed in pigeons, where mixed-action opioids nalbuphine and nalorphine elicited fentanyl-responding (20-45%) in pigeons trained to discriminate a high-dose fentanyl stimulus from saline (Picker et al. 1993). As was highlighted in the aforementioned study, the training dose we used (and the dose used by Bender et al. 2025) was a relatively high dose of fentanyl and thus appreciable levels of substitution may be biased towards drugs with an intrinsic efficacy that is comparable to fentanyl. Our observations with xylazine are more consistent with what would be expected from mixed-action opioids, in that the average appeared to be a partial substitution for fentanyl, where some animals exhibited complete substitution for the training drug and others exhibited no substitution. If xylazine were a low efficacy mu opioid receptor (MOR) agonist, then full substitution would be expected in animals trained to discriminate lower doses of fentanyl. When animals trained to discriminate 0.02 mg/kg fentanyl were probed with 1 mg/kg xylazine, individual animals responded on the fentanyl-lever like they did 10 minutes after they were treated with 0.02 mg/kg fentanyl and 1 mg/kg xylazine (**Fig. 3c**), which is more suggestive that individual differences are underlying the observed substitution rather than low intrinsic efficacy at MORs, or that lower training doses of fentanyl are needed to observe appreciable substitution across subjects. Regardless, lisuride partially substituted (<80%) for xylazine in rats trained to discriminate xylazine from saline (Colpaert and Janssen 1985), and xylazine was recently found to have activity at KORs and D2 dopamine receptors (Bedard et al. 2024), thus irrespective of whether xylazine has low intrinsic activity at MORs or not, it is clear that xylazine seems to be less selective than previously thought.

In anecdotal studies of people who use drugs, a small number of participants report positive sentiments towards xylazine and report seeking out xylazine (Reed et al. 2025; Spadaro et al. 2023). These reports seem to be at odds with preclinical models of substance use disorders, since others have reported that xylazine (i.v.) decreases fentanyl and heroin self-administration in rats (Hochstetler et al. 2025; Khatri et al. 2024), that xylazine has no effect on fentanyl reinforcement and decreases motivation for consuming heroin in rats undergoing progressive ratio tests (Hochstetler et al. 2025; St Onge et al. 2024), and that addition of xylazine to fentanyl was not different from fentanyl conditioned place preference in male Swiss Webster mice (Acosta-Mares et al. 2023). The considerable interanimal variability we observed in the effects of xylazine on locomotor activity, on body temperature, and on the discriminative stimulus effects of fentanyl reveals a possible explanation for the disparate results in human studies and animal models: the pooling of all subjects. Human subjects have remarked on the profound sedative effects of xylazine and how it is undesirable because being knocked out is a “waste of money” and because their belongings are at risk of being burglarized while they are unconscious (Friedman et al. 2022; Reed et al. 2022). Perhaps performing an acute locomotion pre-test with a moderate dose of xylazine (ex: 1-1.78 mg/kg, i.p.) would allow for animals to be separated into groups with low, intermediate, and high sensitivity to the sedative effects of xylazine before initiating preclinical substance use paradigms. Animals that are particularly sensitive to the rate-suppressant effects of xylazine may be reflective of human subjects that experience profound sedation and thereby dislike fentanyl adulterated with xylazine, and pooling these subjects with animals that are not sedated by xylazine may partially explain why rodent studies have reported that xylazine has no effect on or even decreases fentanyl intake.

Xylazine has been commonly reported to exacerbate the lethality of fentanyl and to lead to naloxone-resistant overdose, despite a dearth of preclinical evidence. Although xylazine has been demonstrated to increase the lethality of fentanyl (Acosta-Mares et al. 2023; Smith et al. 2023), naloxone has been shown to precipitate withdrawal in C57BL/6J mice treated with xylazine or fentanyl and xylazine (Bedard et al. 2024), and to transiently reverse fentanyl-induced brain hypoxia (Choi et al. 2024). We sought to further characterize the effects of xylazine on opioid-induced respiratory depression, the most lethal effect of opioids, to further contribute to preclinical efforts to evaluate the reversibility of fentanyl-xylazine respiratory interactions. We found that xylazine alone had transient effects on minute volume, and that a higher dose of xylazine led to a delayed increase in tidal volume (**Fig. 5a**). When a higher dose of xylazine was co-administered with morphine or fentanyl, we observed an enhancement of the respiratory depressant effects of opioids (**Fig. 5b**, **5c** and **6a**). Naloxone reversed respiratory depression induced by morphine and xylazine, but its effect was transient (**Fig. 6a**, **6b** and **6c**), whereas atipamezole or naloxone and atipamezole produced a prolonged reversal of morphine and xylazine’s combined respiratory depressant effects (**Fig. 6b** and **6c**). We then tested morphine and fentanyl alone to assess whether atipamezole could block or reverse opioid-induced respiratory depression. When fentanyl and atipamezole were administered concurrently, it appeared that atipamezole blocked fentanyl-induced respiratory depression (**Fig. 6d**) and may have had a transient reversal effect when administered 30 min after fentanyl. Atipamezole did not appear to have any effect on morphine-induced respiratory depression (**Fig. 6d**), though it may be that 10 mg/kg morphine was not an optimal dose to evaluate this question, since there was no dramatic decrease in respiratory frequency across subjects.

Xylazine has been reported to cause strong, prolonged decreases in brain oxygenation, and increases the duration of fentanyl’s hypoxic effects in male Long Evans rats (Choi et al. 2023). That is when fentanyl is administered alone the initial rapid decrease in brain oxygenation leads to a secondary rebound phase, which is blocked when xylazine is administered alongside fentanyl. Naloxone pretreatment partially decreases fentanyl-xylazine induced brain hypoxia (i.e. the initial hypoxic phase), but only naloxone and atipamezole co-administration was able to counteract the effects of xylazine on the secondary phase of oxygen response (Choi et al. 2024). The delayed increase in tidal volume that we observed following xylazine administration would be consistent with a compensatory response to maintain sufficient oxygenation (**Fig. 5a**). When fentanyl and xylazine were co-administered we observed a significant decrease in respiratory frequency that coincided with a significant increase in tidal volume, but since minute volume was not significantly different from animals treated with fentanyl alone, it appears that a compensatory effect was able to mitigate the added effects of xylazine (**Fig. 5c**). Had we tested higher doses of fentanyl and xylazine we may have recapitulated the findings of Choi et al. where xylazine overwhelms the compensatory second phase of the oxygen response after fentanyl administration. However, we did not continue to test higher doses in our plethysmography tests because our aim was to induce significant respiratory depression but to an extent that was safe (i.e. sublethal). Another plethysmography study found that clonidine decreased minute volume in fentanyl-treated Sprague Dawley rats, and that yohimbine partially reversed the observed decrease, suggesting α2AR activation may be involved in fentanyl-induced respiratory depression (Shaykin et al. 2024). Our finding that atipamezole co-administration blocked fentanyl-induced depression in respiratory frequency, and that subsequent treatment with atipamezole may transiently reverse fentanyl’s effects on respiratory frequency lends further support to the possible involvement of α2AR in mediating the effects of fentanyl on respiration. Our plethysmography results are part of a growing body of evidence that opioid-xylazine overdose is not resistant to naloxone reversal, but that opioid and α2AR antagonists in combination would be more effective at reversing the respiratory effects of fentanyl and xylazine. Although the reversal effects of naloxone are transient when administered after opioid-xylazine treatment, the same is true in the case of fentanyl overdose without xylazine, where Narcan may need to be administered several times to effectively reverse respiratory depression (Pergolizzi et al. 2021; Rzasa Lynn and Galinkin 2018).

α2AR agonists have long been demonstrated to potentiate the antinociceptive effects of opioids (Cakir et al. 2025; Chabot-Dore et al. 2015a; Chabot-Dore et al. 2015b; Spaulding et al. 1979), and several mechanisms for opioid-adrenergic interactions have been identified. MOR agonists and antagonists may bind the extracellular loops of adrenergic receptors, while adrenergic receptor agonists may bind the extracellular loops of MORs (Root-Bernstein and Dillon 2014; Root-Bernstein et al. 2018), perhaps due to shared evolutionary history (Root-Bernstein and Churchill 2021). MORs and α2ARs have been shown to colocalize and to form heterodimers, where activation of only one receptor subtype is sufficient for signaling (Jordan et al. 2003; Vilardaga et al. 2008). Mutations in MORs and delta opioid receptors that have been found in the general public are reported to affect the stability of mu opioid-delta opioid receptor heterodimers (Wu et al. 2021), therefore it is possible that genetic mutations in MORs and α2ARs may affect heterodimerization and may underlie differences in preference or aversion towards xylazine adulteration of fentanyl. Opioid receptor heterodimers may serve as drug targets for future therapeutics (Zhang et al. 2020), and may be targeted using heterobivalent ligands of opioid receptors (Hovah and Holzgrabe 2024; Rehrauer and Cunningham 2023). Future studies may determine which pharmacological effects of opioid-xylazine mixtures are mediated by MORs, α2ARs, MOR-α2AR heterodimers, other receptor interactions (such as imidazoline receptors), or overlapping downstream mechanisms.

In summary, we found that xylazine overall depresses locomotor activity, decreases body temperature, and exacerbates opioid-induced respiratory depression. However, some individual animals were markedly less sensitive to these effects, and we observed considerable interanimal variability despite studying only a small number of male rats. Xylazine partially or even fully substituted for fentanyl in some animals that were trained to discriminate fentanyl from saline but had no fentanyl-like subjective effects in other subjects. Animals trained to discriminate xylazine from saline responded primarily on the saline-lever at doses both above and below the training dose, suggesting the subjective effects of xylazine are variable, dose dependent, and perhaps that these effects are mediated by more than just α2ARs. Naloxone reversal of the combined respiratory effects of morphine and xylazine was transient, whereas the addition of atipamezole produced a long-lasting reversal of respiratory depression. Interestingly, when fentanyl and atipamezole were administered at the same time, atipamezole blocked the respiratory depressant effects of fentanyl, lending further support to the idea that α2AR activation is involved in fentanyl-induced respiratory depression. Altogether, these results highlight that individual differences in the behavioral effects of xylazine and opioid-xylazine mixtures may be a heretofore overlooked aspect of xylazine pharmacology, one that might explain both the polarized opinions of people who use drugs and the counterintuitive preclinical findings that have been reported in rodent models of substance abuse thus far. Furthermore, this work reiterates other preclinical studies that have reported that opioid-xylazine mixtures are not “naloxone resistant”, but that the addition of an α2AR antagonist may be more effective at reversing the respiratory depressant effects of fentanyl whether or not it is adulterated with xylazine.

## Acknowledgments

This work was supported by the National Institute on Drug Abuse of the National Institutes of Health (Award R01DA047967). The content is solely the responsibility of the authors and does not necessarily represent the official views of the National Institutes of Health.

## Author Contributions

K.L.W. and J-X.L. designed the research study. K.L.W. conducted behavioral experiments and performed data analysis. K.L.W. and J-X.L. wrote the manuscript.

---

**Supplemental Fig. 1.**
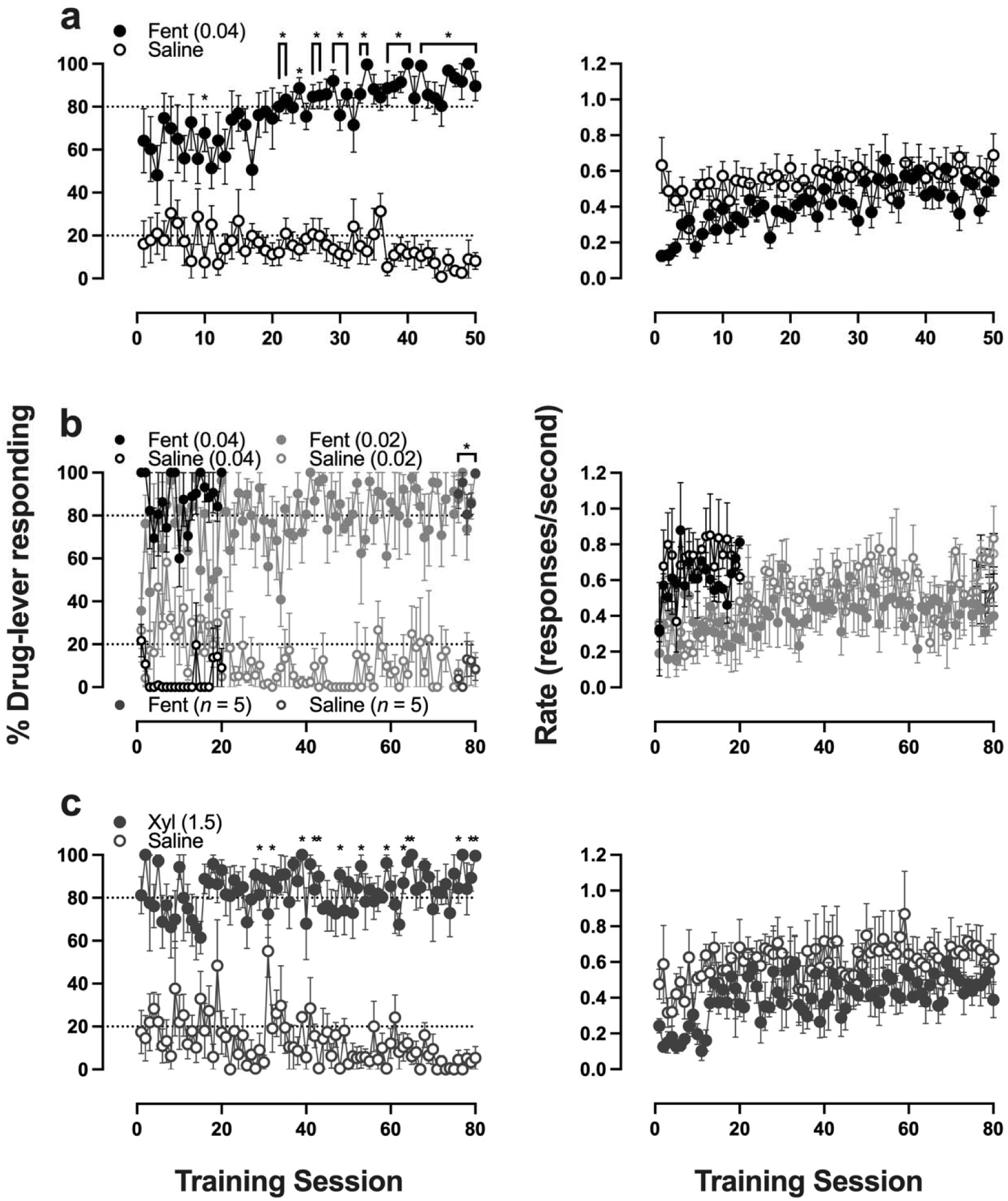
Acquisition curves of the discriminative stimulus effects of fentanyl or xylazine from vehicle in rats. Percent responding on the drug-appropriate lever (*left*) and response rate (*right*) from the first active period during training sessions in rats trained to discriminate 0.04 mg/kg fentanyl (**a**), 0.02 mg/kg fentanyl (**b**), and 1.5 mg/kg xylazine (**c**). Asterisks indicate significant difference from saline sessions.

